# Non-production vegetation has a positive effect on ecological processes in agroecosystems

**DOI:** 10.1101/624635

**Authors:** Bradley S. Case, Jennifer L. Pannell, Margaret C. Stanley, David A. Norton, Anoek Brugman, Matt Funaki, Chloé Mathieu, Cao Songling, Febyana Suryaningrum, Hannah L. Buckley

**Affiliations:** School of Science, Auckland University of Technology, Auckland, New Zealand; School of Biological Sciences, University of Auckland, Auckland, New Zealand; Te Kura Ngahere/School of Forestry, University of Canterbury, Christchurch, New Zealand; HAS University of Applied Sciences, ‘s-Hertogenbosch, The Netherlands; Sichuan Agricultural University, College of Agronomy, Chengdu, China

**Keywords:** Agriculture, biodiversity, non-production vegetation, connectivity, ecosystem disservice, ecosystem function, ecosystem process, ecosystem service, landscape, multi-functionality, resilience, sustainable agroecosystems

## Abstract

An ever-expanding human population, ongoing global climatic changes, and the spread of intensive farming practices is putting increasing pressure on agroecosystems and the inherent biodiversity they contain. Non-production vegetation elements, such as woody patches, riparian margins, and inter-crop and restoration plantings, are vital for conserving biodiversity in agroecosystems and are therefore considered key to sustaining the biotic and abiotic processes underpinning sustainable and resilient agroecosystems. Despite this critical role, there is a surprising lack of synthesis of which types of non-production vegetation elements drive and/or support ecological processes and the mechanisms by which this occurs. Using a systematic, quantitative literature review of 342 articles, we asked: what are the effects of non-production vegetation elements on agroecosystem processes and how are these processes measured within global agroecosystems? Our literature search focussed on the effects of non-production vegetation related to faunal, weed, disease, and abiotic processes. The majority (61%) of studies showed positive effects on ecological processes: non-production vegetation increased the presence, level or rate of the studied process. However, rather than directly measuring ecosystem processes, 83% of studies inferred processes using proxies for ecosystem function, such as biodiversity and soil physicochemical properties. Studies that directly measured non-production vegetation effects focussed on a limited number of vegetation effects including comparisons of vegetation types, farm-scale configuration, and proximity to vegetation. Moreover, studies directly measuring ecosystem processes were similarly limited, dominated by invertebrate biocontrol, predator and natural enemy spillover, animal movement, and ecosystem cycling. We identify research gaps and present a pathway for future research in understanding the ecosystem components and processes that build resilient, sustainable agroecosystems.

## Introduction

Agroecosystems, comprising over 40% of the Earth’s terrestrial surface, are under increasing pressure as the demand for secure sources of food and other resources increases, the increased uptake of intensive agricultural practices, and as climatic processes, disturbance regimes, and habitats continue to be disrupted or degraded (e.g., Godfray et al. 2010, Tilman et al. 2011, Smith et al. 2016). Such pressures are drastically altering agroecosystems and their abilities to provide and sustain services at the accelerating rate of human demand (Rockström et al. 2017). A consequence of this reality is dramatic losses of biodiversity across agricultural landscapes globally (Newbold et al. 2015). There have been discussions whether the biodiversity crisis could be best mitigated via larger conservation set-asides (‘land sparing’), or whether the focus should be on maintaining, retaining, and restoring components of the original diversity of taxa, habitats and ecosystem attributes within the production matrix itself (‘land sharing’) (e.g., Green et al. 2005). The outcome of these discussions is that perhaps both approaches are essential (Kremen 2015), and that even small biodiversity elements within intensive agricultural landscapes are critical for creating **agroecosystems** that exhibit ‘**multi-functionality**’ (Barral et al. 2015; see Box 1 - glossary).

There is increasing evidence (e.g. Tilman and Downing 1994, Srivastava and Vellend 2005, Cardinale et al. 2006, van der Plas 2019), that a positive relationship exists between **biodiversity** and **ecosystem function**, although the exact nature and strength of the relationship is process- and context-dependent (Gamfeldt and Roger 2017, van der Plas 2019). Thus, biodiversity itself is fundamental to sustaining the biotic and abiotic **ecosystem processes** that underpin **sustainable and resilient** agroecosystems (‘**functional biodiversity**’, *sensu* Moonen and Barberi 2008, Martin et al. 2019) and the downstream delivery of **ecosystem services**. There are two contributions of functional biodiversity that must be considered. First, key taxa can have disproportionate effects within agroecosystems, and their loss from a local species pool can have detrimental impacts on biodiversity and ecological processes (Chapin et al. 1997); the decline in bee populations in some regions provides a poignant example of this, having deleterious consequences for pollination and, ultimately, crop productivity (Vanbergen and the Insect Pollinators Initiative 2013). This highlights that, while species richness is important, more critical is the functional roles that particular species play in agroecosystems, such as in multi-trophic processes (Soliveres et al. 2016) and in the decomposition of litter and detritus (Gessner et al. 2010).

Second, **non-production vegetation** elements provide the surrounding context in which the production components of agroecosystems are embedded. These non-production vegetation elements vary in their size, composition and spatial arrangement in agricultural landscapes; these aspects thus determine the types of processes that can be supported in agroecosystems. A number of government programmes across the globe, such as agri-environment schemes in the UK, USA, and Europe, have focused on creating/maintaining non-production vegetation components in agricultural landscapes, with variable outcomes (e.g., Batáry et al. 2015, Wood et al. 2015, Zmihorski et al. 2016, Jones et al. 2017). Multiple processes are required to achieve a resilient ecosystem (Oliver et al. 2015), but we lack synthesis in our understanding of the role of non-production vegetation elements in multiple ecosystem processes. If biodiversity positively affects ecosystem function, we need to understand which and how non-production vegetation elements support ecosystem processes.

In this review we evaluate the role of non-production vegetation elements within agricultural landscapes, in terms of supporting both abiotic and biotic processes at different spatio-temporal scales, and thus their importance in achieving sustainable, resilient agroecosystems. Such non-production vegetation includes linear features like hedgerows, shelterbelts, corridors, and riparian buffers, as well as patches of vegetation composed of restoration plantings and remnant patches, and other inter-crop components such as underplantings, buffer strips, and cover crops. In particular, we ask: What functions and processes are associated with non-production vegetation components and how are these processes measured within global agroecosystems? We used a quantitative review methodology (Pickering & Byrne, 2014) to search the relevant literature to obtain a sample of research articles using keywords. We scored papers based on how non-production elements were measured, and the contributions to biodiversity and ecosystem function(s) demonstrated by these non-production elements. We use these results to make recommendations for future research that will support the maintenance and enhancement of the world’s sustainable and resilient agroecosystems.

### Quantitative literature review

We searched the international literature for relevant articles following a modified systematic review protocol (Pickering & Byrne 2014). Relevant articles were defined as those that described primary research conducted on farmland that addressed questions regarding any ecological function(s) associated with non-crop vegetation. Five distinct search strings, comprised of various keywords (Table S1) were designed to find relevant papers addressing the following broad topics related to non-production vegetation in agroecosystems; faunal diversity and use of vegetation; spatial arrangement of vegetation in the landscape; pests and disease in vegetation; weeds in vegetation; abiotic functional responses to vegetation. The hit rate of 30 scholarly databases was tested using all five search strings, and nine were selected based on the proportion of relevant hits: BASE, BioOne, Google Scholar, JSTOR, Jurn, ProQuest, Science Direct, Scopus, and Web of Science. Each search string was entered into each database in turn, and all relevant articles were downloaded (where relevance was determined by reading the abstract). If no relevant articles were found for 100 hits, we moved on to the next search. For every relevant article found, we also checked all citing articles using the “Cited by” function in Google Scholar. After the initial two databases were searched, we extracted all keywords from papers and ranked them by the number of occurrences. We updated the search strings to include any commonly-used keywords that were missing from the strings (Supplementary Materials Table S1). Finally, once all searches were complete, we extracted and cross-referenced all reference lists from the downloaded articles using ParsCit (Kan et al. 2011), and checked all papers that were cited three or more times and were not already in our collection.

A total of 704 articles were read and 342 were included using the criteria that they were: (1) empirical (not modelling or meta-analysis) studies within agroecosystems; and (2) they at least discussed the effects of non-production vegetation on processes, not just biodiversity, within agroecosystems. A range of initial variables, were extracted from every included article, describing the study design, the stated aims/questions of the study, the taxa studied (e.g., birds, invertebrates, mammals), the non-crop vegetation type (e.g., woody, herbaceous) and configuration (e.g., forest fragments, hedgerows, corridors) (Table S2), the spatial grain and extent of the study (e.g., field margin, field, farm, catchment, region). We noted whether the inferences made in the paper were regarding ecological processes, biodiversity or both, and what processes were studied, such as biocontrol, pollination, nutrient cycling (Table S3). Finally, we recorded the methods that were used to measure processes directly, including direct observations of animal movements or feeding events, herbivory rate observations, respiration or decomposition rates, or whether the process was inferred via indirect observations or measurements, such as using relative differences in predator relative abundance to infer predation rates between habitat types or using soil physicochemical variables to infer nutrient cycling.

The 342 relevant research articles had a large global distribution (Fig. S1), and resulted in 229 described independent studies about fauna, 61 studies about soil and water processes, 32 studies about weeds, and 17 about diseases (human diseases, n = 5; other animal diseases, n = 2; plant diseases, n = 10); note that a few articles included more than one independent study. A total of 500 different research questions were asked within the articles regarding the effects of non-production vegetation on agroecosystem processes. Globally,

### Non-production vegetation positively affects agroecosystem function

Our synthesis shows that the study aims, as stated by reviewed articles, address a wide variety of agroecosystem processes (Fig. 1). Across the 342 reviewed articles and 500 independent research questions posed within these articles, the majority of effects of non-production vegetation on ecological processes were positive in that non-crop vegetation increased the presence, level or rate of the studied process (58%; n = 290). Of these processes, 10% (n = 51) would be classified as ‘ecosystem disservices’ because they contribute negatively to human-preferred outcomes in agroecosystems. Of the remaining 90% (n = 449) of processes tested, non-crop vegetation caused increases in 61% of the processes (faunal biodiversity, n = 146; faunal processes, n = 64; soil and water processes, n = 53; weed processes, n = 4; disease processes, n = 5), caused decreases in 4% of the processes, caused variable responses (where the outcome depended on another factor such as vegetation type, taxon/taxa involved, landscape configuration or season) in 14% of the processes and was non-significant or unclear in 21% of processes. Of the ‘ecosystem disservice’ processes, non-crop vegetation improved (i.e., reduced the disservice) 35% of all studied processes (faunal processes, n = 11; weed processes, n = 5; diseases processes, n = 2).

**Figure 1:**
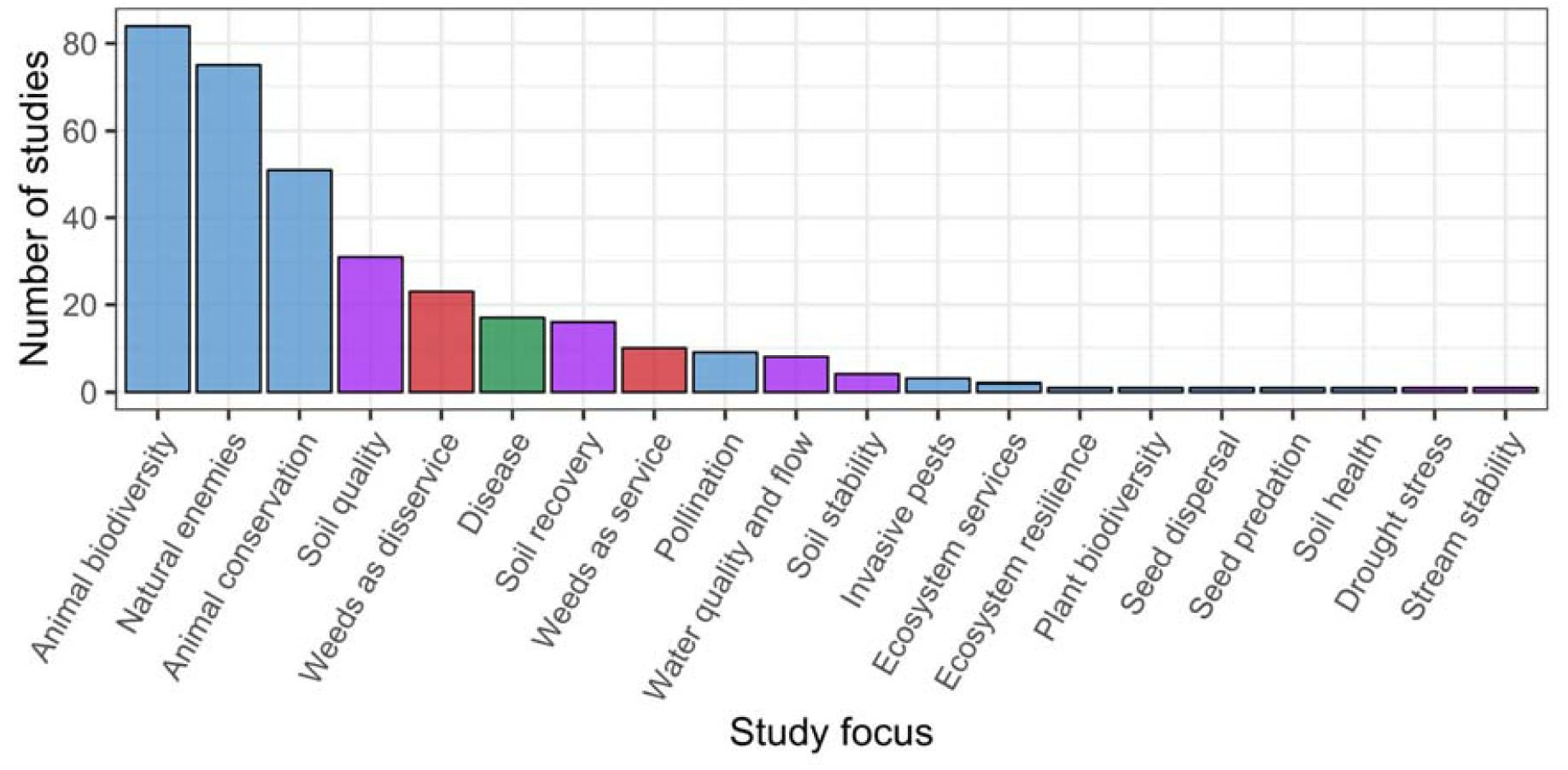
The number of total reviewed studies investigating different study aims. Over 90% of the total 342 reviewed studies focused on seven broad study aims with respect to the effects of non-production vegetation: animal biodiversity, natural enemies (i.e., invertebrate biocontrol agents), animal conservation, soil quality, weeds as an ecosystem disservice, disease occurrence and/ or spread in agroecosystems (animal, plant and human), and soil recovery, with the remaining <10% of reviewed papers focusing on an additional thirteen study aims. Blue bars represent fauna-related processes, purple bars represent soil and water-related processes, red bars represent weed-related processes and the green bar represents disease-related processes.

The most-commonly studied types of non-production vegetation elements were classed as ‘patches’, meaning any non-linear fragment that was either planted, or more frequently remnant in the agricultural landscape (n = 183; Fig. 2). Studies that tested at least two different types of non-production vegetation elements (‘mixed’; n = 109) and hedgerows or field margin plantings (‘borders’; n = 98) were also studied more frequently than any of the other types, surprisingly including corridors (n = 5; Fig. 2), which have been touted as important connecting elements in agriculture landscape (Correa Ayram et al. 2016). Whether or not non-production vegetation was present in the landscape (‘provision’; n = 103) and the comparison of different non-production vegetation types (n = 95) were the most commonly tested effects of non-production vegetation on agroecosystem processes. Hedgerows/borders and shelterbelts showed relatively more increasing (rather than variable, decreasing or non-significant) effects than other vegetation types, across a range of tested effects (Fig. 2).

**Figure 2:**
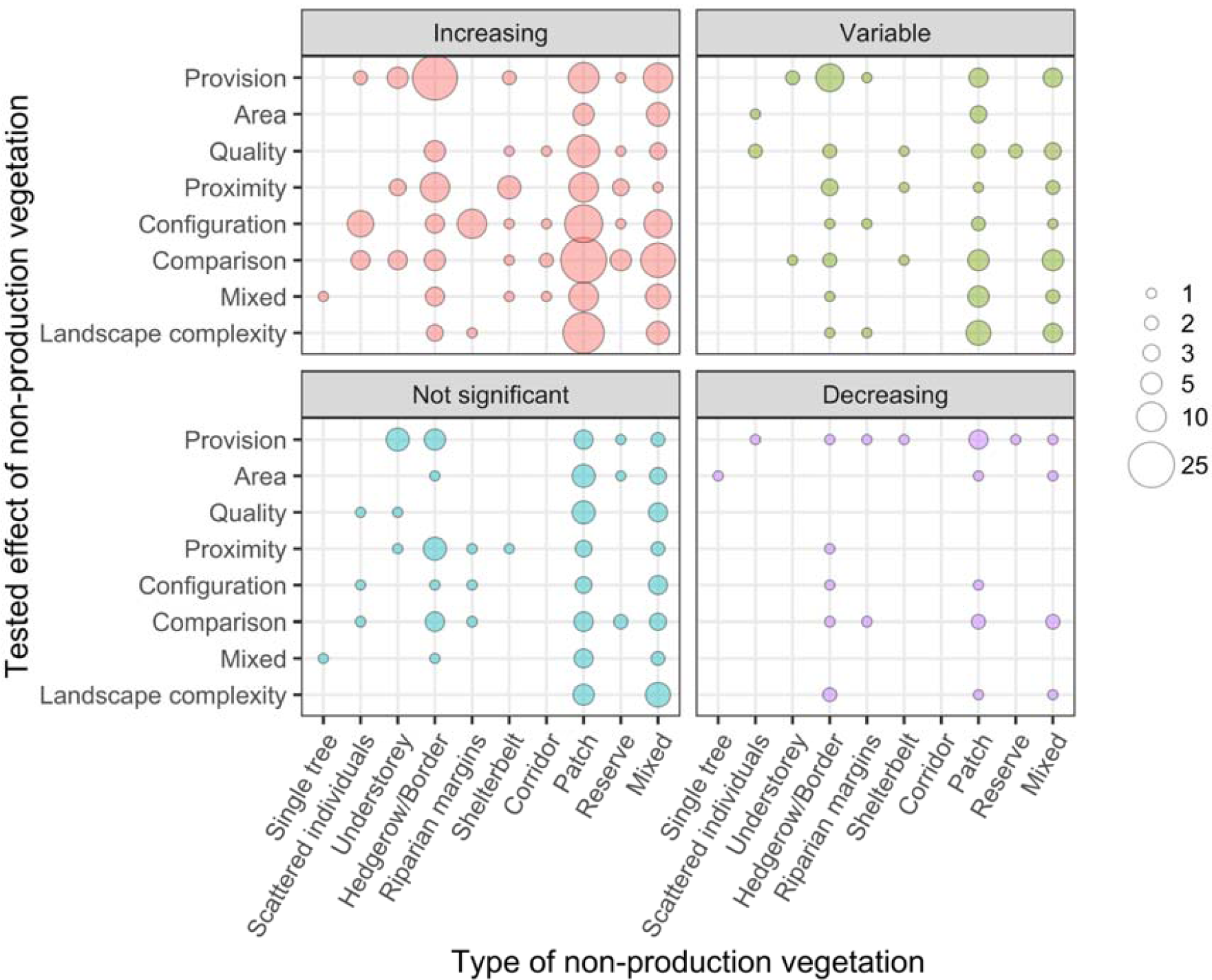
The number of studies grouped by the type of non-production vegetation and its tested effect, as stated in each of the 500 research questions posed by the 342 reviewed articles, including those both inferring and directly-measuring processes. The outcomes of these tests were classified as ‘increasing’ (n = 287; ecosystem processes that were significantly improved by the effect non-production vegetation; in the case of ‘disservices’, the process was lessened or prevented), ‘decreasing’ (n = 26; processes that were lessened or prevented by the effect of non-production vegetation; in the case of ‘disservices’, the process was increased or enhanced), ‘not significant’ (either a non-significant [n = 74] or unclear [n = 26] effect) and ‘variable’ (n = 83; where the outcome depended on another factor such as vegetation type, taxon/taxa involved, landscape configuration or season).

### Agroecosystem processes are mostly inferred, not directly measured

The ability to draw strong conclusions regarding the effects of non-production vegetation on agroecosystem processes is dependent on the degree to which processes are measured directly compared to whether indicator or proxy variables are used to represent those processes. In our review, very few studies directly measured the effects of non-production vegetation on agroecosystem processes. The majority of the 500 research questions from the 342 reviewed articles (83%; n = 416) were posed using variables that have only been hypothesised to represent ecosystem function, such as biodiversity and soil physicochemical properties. There is only variable and/ or weak evidence for many hypothesised causal relationships between such proxy variables and ecosystem functions, such as the links between biodiversity and soil carbon, decomposition rates, or herbivory (van der Plas 2019) or the links between indicators of soil properties and processes such as nutrient cycling and water quality (Bünemann et al. 2018). In other cases, links have been shown to be weak or absent. For example, land use types are often used as a proxy for ecosystem function, but this has been shown to be unreliable (Bünemann et al. 2018) and there have been suggestions that more work needs to be done to model soil processes before causal relationships can be determined between processes and easy-to-measure indicator or proxy variables (Vereecken et al. 2016). Thus, our results show that greater research emphasis needs to be put on directly measuring ecosystem functions in agroecosystems associated with non-production vegetation and disentangling their relationships with other variables such as biodiversity and abiotic properties.

The research questions that used direct measurements of ecosystem processes rather than inferred links (n = 84 out of 500) examined a limited range of taxa and ecosystem processes and used a limited set of methods to measure those processes (Table 1), revealing significant research gaps in this field. For example, 43% of studies on ecosystem processes related to fauna in non-production vegetation elements, looked at invertebrate biocontrol and natural enemies of crop pests, such as parasites (n = 26 out of 61 studies); indeed, the majority of these were invertebrate-related studies (n = 38), while the remainder studied birds (n = 11), mammals (n = 8), multiple taxa (n = 3), and reptiles (n = 1). For soil and water-related ecosystem processes, 19 out of 20 studies looked at ecosystem cycling processes such as decomposition, carbon cycling and nutrient cycling; no studies directly measured soil erosion, a key process in agroecosystems often touted to be positively affected by non-production vegetation. Only three weed studies directly measured processes by monitoring weed invasion over time; all other studies, including 30 on weeds and 17 on diseases inferred processes sampling the presence and/or abundance of weed- and disease-related taxa in different non-production vegetation elements and the production matrix.

**Table 1:** Number of research questions directly testing different agroecosystem processes and the methods used to measure each process type. Ecosystem cycling refers to measurements of the flux and cycling of water, carbon or nutrients. Habitat selection refers to measurements of mobile animals that moved into (or, in the case of weeds, were excluded from) non-production vegetation elements. Spillover refers to the movement of animals out of non-production vegetation elements. Matrix invasion refers to the spread of weeds into non-production vegetation elements. Direct observation refers to methods that monitored animal movement such as GPS trackers and capture-recapture studies. Soil emission measurements refers to studies that used experimental methods, such as incubation, to record respiration and gas exchange of soils.

| Study type | Process measured | Method | N |
| --- | --- | --- | --- |
| Soil and water | Ecosystem cycling | Litter bag experiments | 6 |
| Soil and water | Ecosystem cycling | Soil emission measurements | 9 |
| Soil and water | Ecosystem cycling | Isotope measurements | 1 |
| Soil and water | Ecosystem cycling | Mass balance measurements | 1 |
| Soil and water | Ecosystem cycling | Multiple methods | 1 |
| Soil and water | Ecosystem cycling | Piezometer experiment | 1 |
| Soil and water | Soil sedimentation | Sediment accumulation | 1 |
| Fauna | Animal movement | Direct observation | 8 |
| Fauna | Animal movement | Population genetics | 2 |
| Fauna | Biocontrol | Food consumption amount or rate | 15 |
| Fauna | Habitat selection | Direct observation | 6 |
| Fauna | Pest animal movement | Direct observation | 1 |
| Fauna | Pest competition | Direct observation | 1 |
| Fauna | Pest habitat selection | Direct observation | 4 |
| Fauna | Pest herbivory | Food consumption amount or rate | 5 |
| Fauna | Pest predation | Direct observation | 1 |
| Fauna | Pest predation | Food consumption amount or rate | 1 |
| Fauna | Predation | Food consumption amount or rate | 4 |
| Fauna | Predation | Direct observation | 2 |
| Fauna | Spillover | Food consumption amount or rate | 9 |
| Fauna | Spillover | Direct observation | 2 |
| Weeds | Habitat provision | Individual counts | 1 |
| Weeds | Matrix invasion | Tree ring counts | 1 |
| Weeds | Matrix invasion | Individual counts | 1 |

Of the research question that addressed directly-measured processes, the majority (79%; n = 66) showed increases in the presence, level or rate of the ecosystem process, compared to few that showed variable (n = 8), decreasing (n = 3) or non-significant (n = 7) effects (Fig. 3). This suggests that, despite the relatively minimal use of direct process measurements, there is quantitative support for the idea that non-production vegetation beneficially affects agroecosystem processes. Across all tested vegetation effects, ecosystem cycling (n = 19), biocontrol (n = 15), spillover (n = 11), and animal movement (n = 10) and were the processes most frequently measured directly, with almost all of these (82 %) showing positive (increasing) outcomes. Certainly, the last two decades has seen a proliferation of studies on the role of non-production vegetation in supporting beneficial invertebrates in agroecosystems, in terms of providing habitat and resources (e.g., Knapp and Řezáč 2015, Saunders et al. 2016), facilitating their movement into adjacent farmland (e.g., Inclán et al. 2015), and for generally enhancing biocontrol of invertebrates that impact on crop production (e.g., Pywell et al. 2015). There were clear positive effects of several different types of non-production vegetation elements, in comparison to production areas, on ecosystem cycling, such as decomposition, soil respiration, and nitrogen mineralisation (Fig. 3). Conversely, our review also revealed gaps in the types of processes investigated. For example, few studies directly measured processes related to pest competition and movement, the dispersal and invasion of weeds into the production matrix, and other abiotic processes, such as soil erosion and sedimentation (Fig. 3). Further, our results suggest that processes related to non-production vegetation were not commonly directly-tested at landscape scales (e.g., landscape complexity effect, Fig. 3), with many effects tending to be inferred indirectly via indices of land use intensity and composition (e.g., Jonsson et al. 2012).

**Figure 3:**
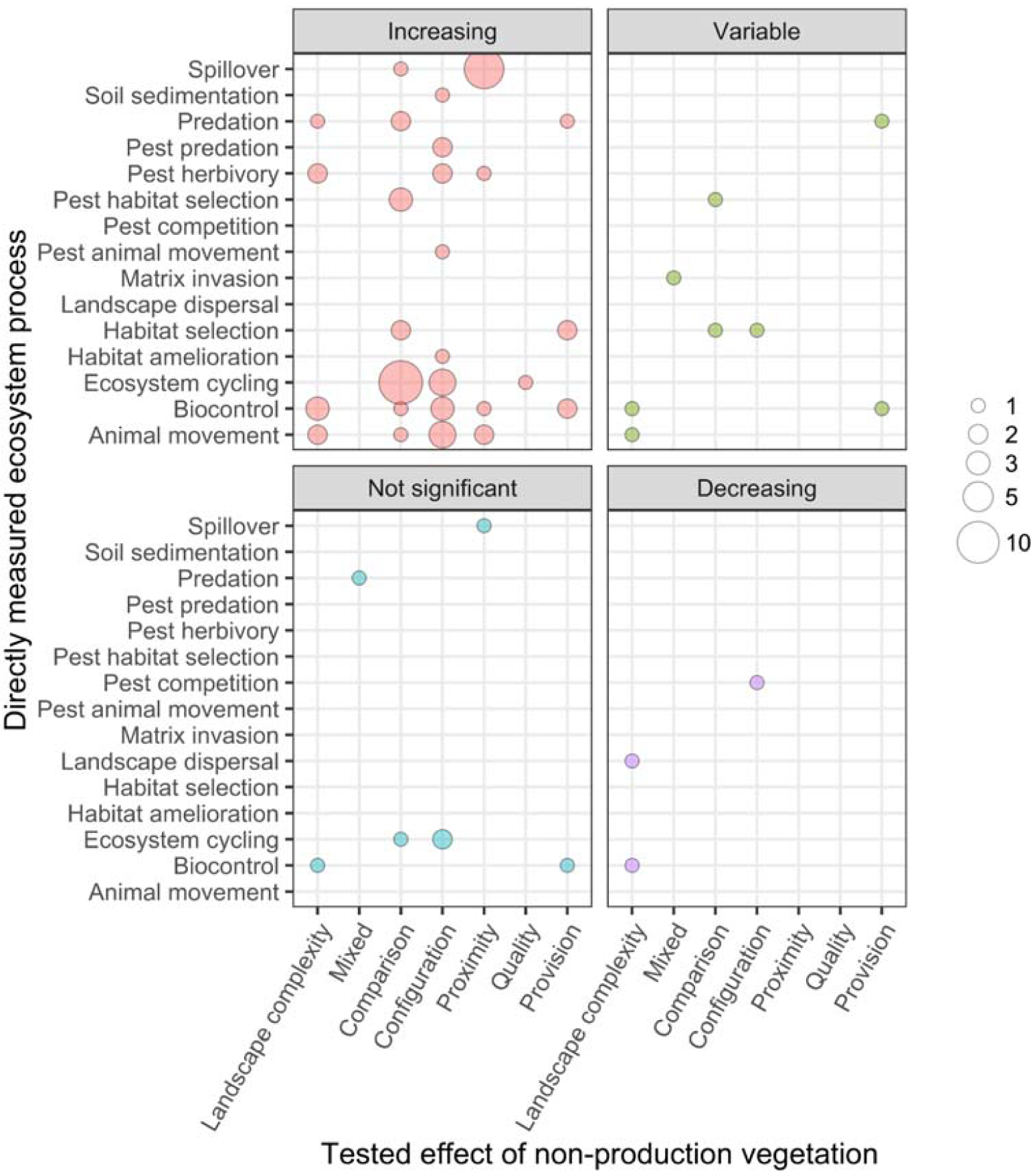
Cross -tabulation results for 84 reviewed studies where processes were directly measured relating the stated, tested effects of non-production vegetation and measured ecosystem processes, grouped by study outcome. As per Figure 2, outcomes of these tests were classified as ‘increasing’, ‘decreasing’, ‘not significant’ or unclear effect and ‘variable’.

### Available methods should be harnessed to improve the direct measurement of processes

Eleven methods were employed across the 84 research questions emerging from studies directly measuring non-production vegetation effects on ecosystem processes (Table 1). Of these 11 methods, only six methods were used in more than one study, and mainly comprised direct observations and food consumption amounts or rates for faunal studies and litter bag experiments and soil flux measurements. This suggests that both the biotic and abiotic methods typically employed are limited to those that are easier to conduct in the field; for instance, vegetation effects on invertebrate predation is likely easier to document than those on competition, and disease transmission and soil gas flux measurements are relatively straightforward compared to those of erosion rates or nutrient and water flow dynamics.

There is a range of underused methods available for measuring ecosystem processes that could be usefully applied in agroecosystem research. For instance, landscape genetics methods, which are used extensively in natural habitats (Manel et al. 2013), could be employed more widely in agroecosystem studies to reconstruct the movement, current gene flow and past history of native fauna in the fragmented habitats of agroecosystems (e.g., Jaffé et al. 2016). Temporal studies that track the movement of native and pest animals, weeds, and diseases through agroecosystems would allow us to assess the role of non-production vegetation elements in providing both connectivity for native species (affecting ecosystem resilience) and pathways for the spread of unwanted organisms. For example, non-invasive and low-cost methods for direct tracking of animals are available such as the use of rubidium as an isotope tracer to track invertebrate movement (Payne and Dunley 2002), or GPS to quantify the movement of mobile macrofauna (e.g. Neilly and Schwarzkopf 2017). Temporal studies involving experimental tracers could also be used to assess how non-production vegetation elements affect ecosystem processes such as sedimentation and run-off (e.g., Mabit et al. 2018), and the potential level of below-ground connectivity between different ecosystem components. Indeed, the roles of non-production vegetation elements for farm-to-catchment scale water flow and movement was particularly understudied in the literature we reviewed.

Experimental studies similarly provide strong inference for tests of process under manipulation; even large-scale manipulative experiments can be logistically easier to conduct in agroecosystems than in native ecosystems (e.g., Resasco et al. 2017). Experimental approaches in agroecosystems could also include subsequent measurements of social and economic outcomes, such as crop yield, meat yield as outputs, social outputs (e.g., Maseyk et al. 2017), thus expanding the scope of the experiment to test for effects of non-production vegetation elements on multi-functionality. Greater natural history and species-level understanding is needed to build knowledge of the processes that support resilient ecosystems (Oliver et al. 2015), and enable the scaling up of key processes using appropriate modelling to predict landscape-level ecosystem outcomes (Padulles Cubino et al. 2018).

### The role of non-production vegetation is embedded within the socio-ecological context

This review has shown that non-production vegetation can support the processes that underpin functional biodiversity in resilient, sustainable agroecosystems. However, fundamentally, agroecosystems are a human creation and so our choices and subsequent behaviours, such as maintaining or restoring non-production vegetation patches in agricultural landscapes, ultimately determine the structure and functioning of these landscapes (Fig. 4A; Landis 2017). Our review shows that there are gaps in our understanding of the broad range of agroecosystem processes that are likely to be affected by non-production vegetation elements. If we aim to achieve sustainable and resilient agroecosystems, research efforts to expand biological knowledge must be embedded in the ‘cultural context’ of agroecosystems, or we risk missing the role of people and the influence of their decisions in maintaining or disrupting the key biological processes and relationships (Fig. 4B).

**Figure 4:**
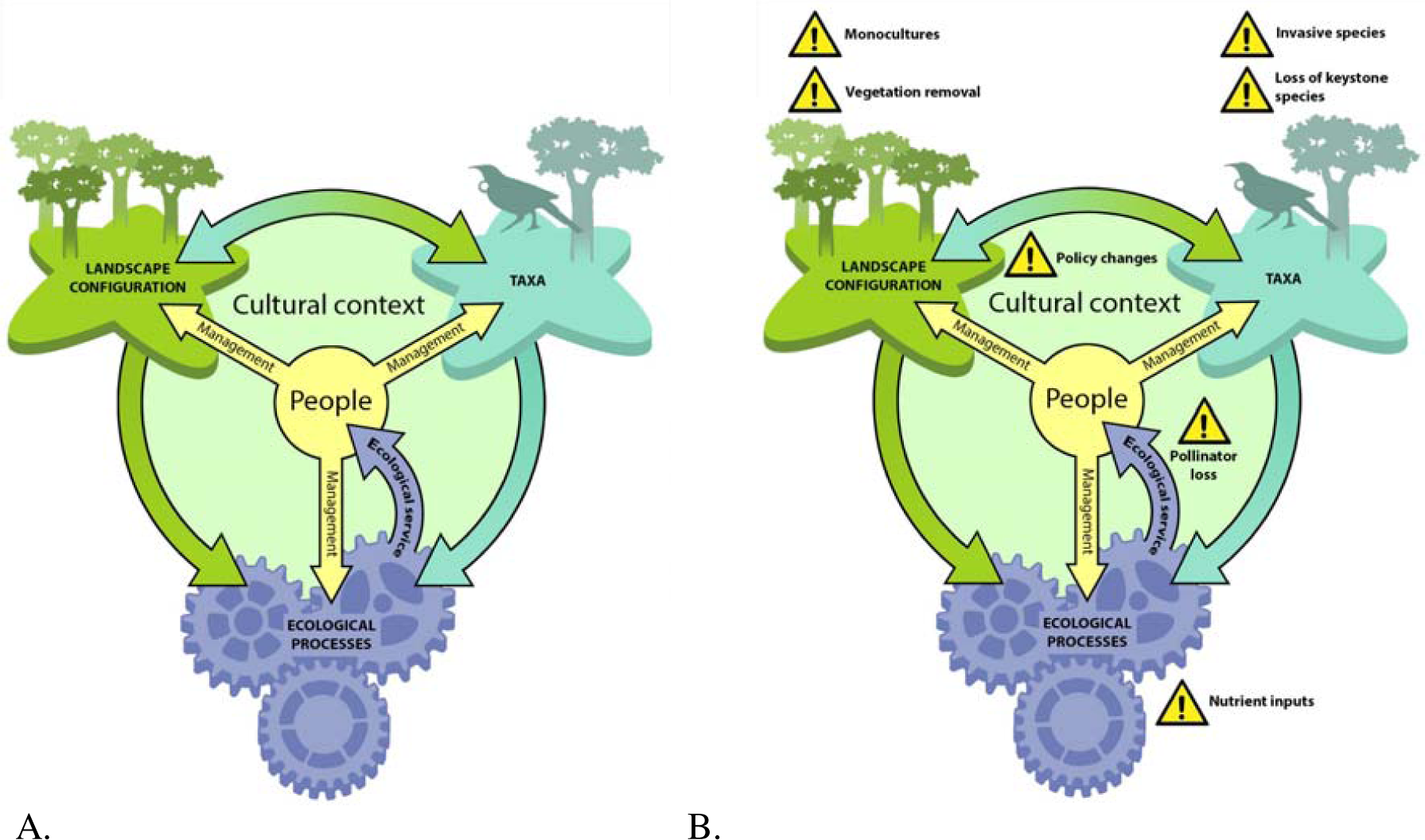
(A) People create agroecosystems and the management of these agroecosystems depends on the cultural context, which underpins landowners’ decision making. This human centre-point drives three components of multi-functionality and ecosystem resilience: (1) the taxa present in the agroecosystem, (2) the landscape configuration (including amount, arrangement and condition) of the non-production vegetation elements in the landscape, and (3) the ecological processes that are operating; taxa and landscape configuration interact to determine the ecological processes. (B) Disruption, due to management decisions, policy changes, and/or human-caused biotic and abiotic alterations, can occur for any of the components and result in the collapse in agroecosystem function and a loss of multi-functionality. To better understand how to maintain or enhance resilience, research needs to be done at multiple scales and focus on under-emphasised processes (see Box 2).

People make land management decisions, including those involving non-production vegetation, for a wide variety of reasons including economic consideration, personal values, and their knowledge of biodiversity and its role in the farming landscape (Norton and Reid 2013). For example, a primary driver of land management decisions is the economics of the farm business, but this is not always in conflict with good functional biodiversity management (Smith and Watson, 2018) and further, can be incentivised by local and national government policy (Hanley et al. 2012). How we value biodiversity intrinsically, such as for the enjoyment of native bird song, the provision of pollination or harvestable material, or for soil nutrient mitigation, has been shown to provide both incentive for good biodiversity management and economic benefits (Cáceres et al. 2015). Traditional ecological knowledge and the cultural importance of particular species or habitats can positively influence the maintenance and enhancement of non-production vegetation and associated biodiversity (e.g., Ruiz-Mallén and Corbera, 2013); considerably more work is required to fully understand the contribution of these decisions to enhancing agroecosystem processes besides maintaining diversity and preventing extinctions. In contrast, fear of native species such as large mammalian predators, or other forms of human wildlife conflict can impact on the management and use of non-production vegetation in farming landscapes (Sitati et al. 2005). Pest or disease vector taxa provide an economic incentive for management and control; where such species also negatively affect native biodiversity, this provides a win-win for farming and good biodiversity management. For instance, the control of the invasive brushtail possum (*Trichosurus vulpecula*) in New Zealand, which is both a vector for bovine tuberculosis infection in cattle and a rapacious predator on native birds, has led to significant economic and ecological gains (Byrom et al. 2016). Conversely, economic returns reaped via the management of non-production vegetation for one ecosystem function (e.g., pollination) can at times occur at the expense of other processes (e.g., biocontrol) (Shackelford et al. 2013). Thus, management decisions in agroecosystems arise from such complexities related to the socio-ecological context and often result in trade-offs between competing viewpoints (Saunders et al. 2016), potentially disrupting one or many parts of the system (e.g., Fig. 4B).

In light of this cultural and ecological context (i.e., land management decisions are at the farm level, processes occur at multiple scales, and management actions need social license), ecological understanding needs to be built within a framework that encompasses these three main components: the configuration of non-production vegetation elements in the landscape, the role of important taxa/functional groups and how these taxa are supported by non-production vegetation, and the interplay of these components with management decisions (Saunders et al. 2016). However, research should not be limited to a narrow set of topics to inform landscape design, but on as many processes as possible (Landis 2017). We advocate for transdisciplinary research on the role of non-production vegetation that capture a broad range of cultural and ecological processes across multiple scales; such research is necessary for informing decision making that will achieve sustainable and resilient systems that support higher functional biodiversity.

Further, because agroecosystem decision making operates at the farm level, while functional biodiversity often operates at coarser scales (Kleijn et al. 2019), multi-level thinking is required to manage functional biodiversity across spatial scales (Box 2). The spatial scale at which ecosystem measurements should be taken needs to consider: (1) the scale of the organisms or the processes under study, (2) the size and arrangement of the non-production vegetation elements to be evaluated, and (3) the logistical or other constraints of the methods used to measure the process. Where possible, pilot studies or prior information from the literature (e.g., Welsch et al. 2019) should be used to justify the spatial scale of sampling. Further, fragmented landscapes cannot sustain all species because some, for instance, require large home ranges; however, where multi-functionality is the goal, rather than solely conservation of individual species, resilient systems need not contain maximal landscape biodiversity to be able to adapt to recover after perturbations such as drought or fire, or adapt to shifting conditions such as increasing temperatures or rising nutrient inputs (Lindenmayer et al. 2008). Instead, the focus should be on restoring, enhancing and manipulating non-production vegetation elements to create connected, structurally complex agricultural landscapes with a diversity of species across key functional groups at multiple scales (Fisher et al. 2006).

## Conclusions

Non-crop vegetation elements in agroecosystems make a significant, positive contribution to biodiversity and ecosystem functioning. Nonetheless, there are gaps in our understanding that can be filled by studies focussing on disentangling the role of different taxa, (or functional groups) across non-production vegetation elements, incorporating a wider variety of processes and spatial scales, and employing novel and underused methods. Indeed, if we are to increase and enhance functional biodiversity, future efforts should be focussed on measuring and monitoring relevant biotic and abiotic ecosystem processes at landscape scales within the context of farm management scenarios. This will lead to new insights into how the types, amounts, and arrangements of non-production vegetation elements, mediated by decision making at the community level, can result in resilient agroecosystems into the future.

## Acknowledgements

This work was supported by funding from the New Zealand Ministry of Business, Innovation and Employment (New Zealand’s Biological Heritage NSC, C09X1501). We thank Mia Jüllig for help with figure creation (http://www.paperdog.co.nz/about/).

## Boxes

### Box 1 Glossary of terms

**Agroecosystems** are ecosystems that have been modified from their natural state through time by farming practices, and may also contain other land uses such as settlements, conservation land or other industry. Like natural ecosystems, agroecosystems are composed of organisms (including humans) interacting with each other within an abiotic (chemical and physical) context.

**Non-production vegetation** elements, which are not directly involved in the farm operation, include trees and woody shrubs, herbaceous vegetation and wetlands. These elements can vary in their size, shape, arrangement in the agricultural landscape, and in their species composition.

These vegetation elements are the main components of ‘**functional biodiversity**’ on farms, in that they support a range of flora and fauna that contribute to key **ecosystem processes**, such as animal dispersal, pollination, and carbon and nutrient cycling, that determine overall **ecosystem function**.

Human-derived benefits may arise from functioning ecosystems in the form of **ecosystem services**, such as clean water, healthy soils, increased crop production, and overall well-being. Conversely, functional biodiversity may lead to ecosystem disservices, such as the facilitation of pest, disease, or weed spread and the knock-on effects for humans.

**Multifunctionality** can arise in agricultural landscapes with high functional biodiversity, intact ecosystem processes, and which are generating multiple ecosystem services.

**Sustainable and resilient** agroecosystems are those which exhibit the ability as a whole to withstand disturbance and/or species loss while supporting farm production into the future.

### Box 2. Filling the gap: A synthetic approach for building understanding of non-production vegetation effects on agroecosystem processes

The main components of resilient agroecosystems are likely to operate at, and interact across, different spatial scales. Therefore, the scale of observation and the methods used differs for different processes and ecosystem components (taxa, configuration of non-production vegetation, agroecosystem processes, and people). Appropriately matching the scales of observation and measurement of processes is critical to deepening our understanding of agroecosystem function.

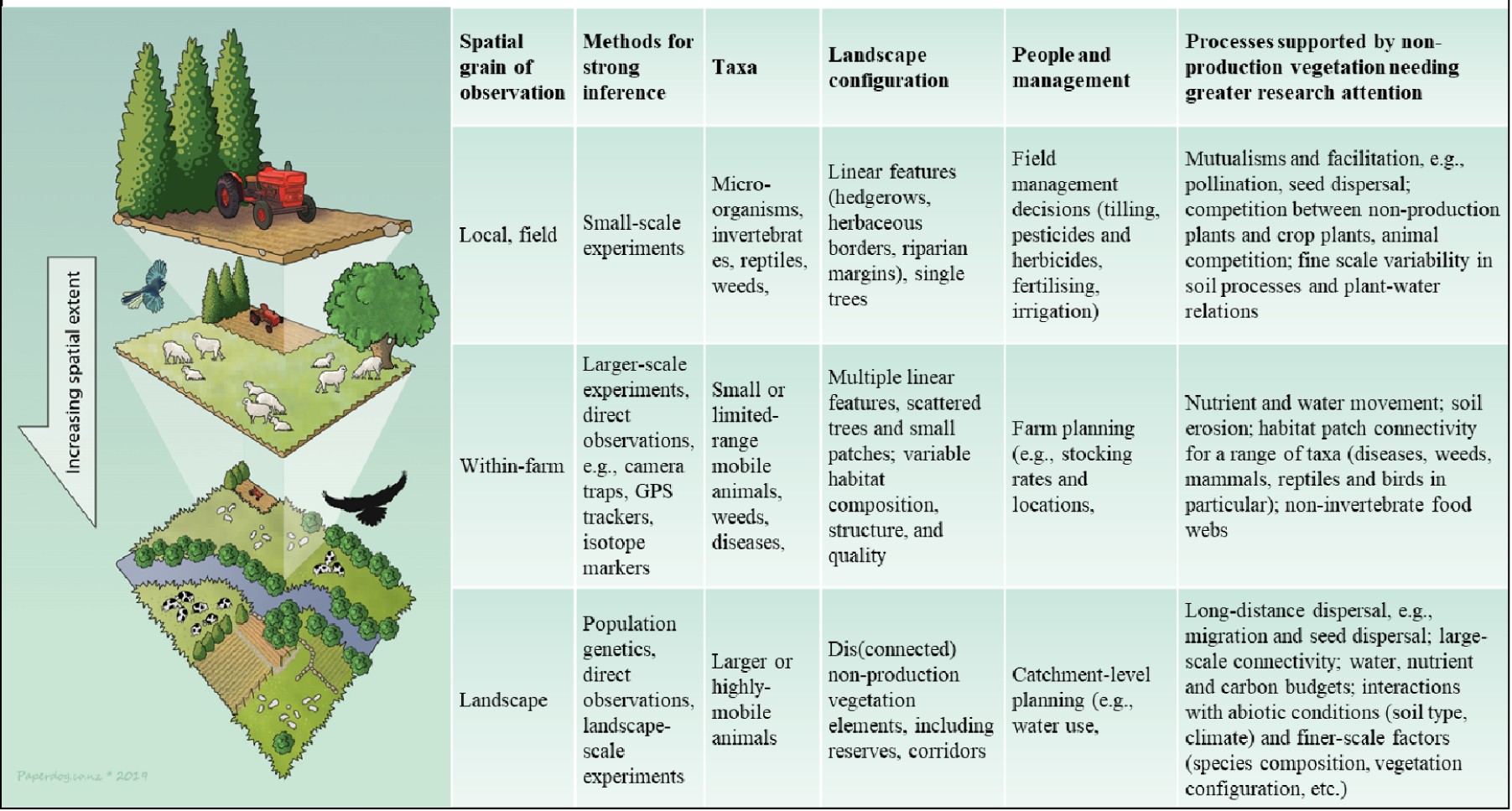

## Supplementary material

**S1 Figure:**
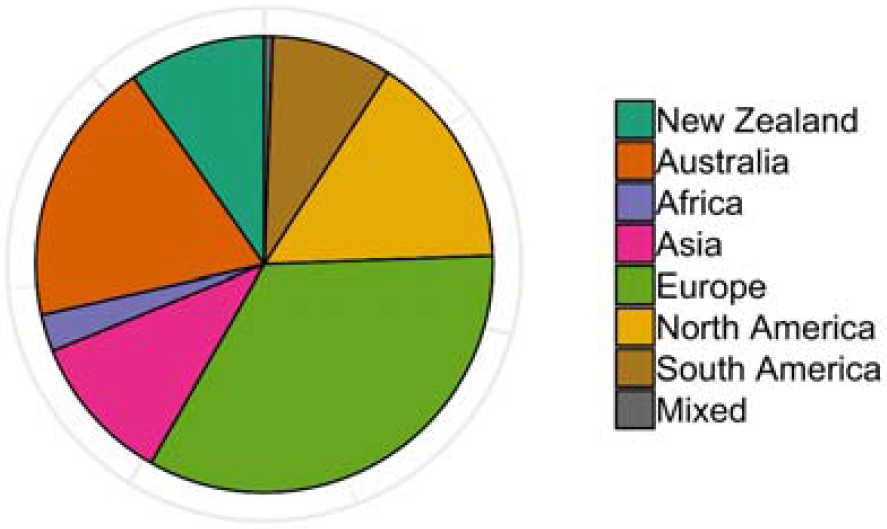
Relative frequency of studies from all searches categorised by country or continent in which the research occurred.

**Table S1:** Search strings used in systematic literature review, for each of the five general themes, both initial strings and after updating after cross-checking against article keywords. Strings formatted for use in Google Scholar; other scholarly databases require different formatting.

| Topic | First Search String | Updated Search String |
| --- | --- | --- |
| Fauna | (native AND habitat) AND (fauna OR animal OR invertebrate OR bird OR bat) AND (biodiversity OR remnant OR restoration OR richness OR corridor OR barrier OR connect OR dispersal OR migration) NOT (fish OR aquatic) | (native AND habitat) AND (fauna OR invertebrate OR bird OR bat) AND (biodiversity OR remnant OR restoration OR corridor OR barrier OR connect OR dispersal OR migration OR recolonization OR revegetation OR occupancy) NOT (marine OR aquatic) |
| Spatial | (biodiversity AND farm) AND (agroecology OR beef OR lamb OR cattle OR sheep OR grazing) AND ("ecosystem services" OR "ecological function" OR indicator OR richness OR scale OR landscape OR patch OR layout) | (biodiversity AND agriculture) AND (agroecology OR beef OR lamb OR cattle OR sheep OR pasture) AND ("ecosystem services" OR "ecological function" OR indicator OR richness OR scale OR landscape OR patch OR layout) |
| Disease | (habitat AND invasive) AND (pest OR alien OR exotic OR disease OR weed OR pathogen OR predator) AND (connectivity OR dispersal OR fragmentation OR corridor) NOT (fish OR aquatic) | (habitat AND invasive) AND (pest OR alien OR exotic OR disease OR weed OR pathogen OR zoonoses OR predator) AND (connectivity OR dispersal OR fragmentation OR corridor OR revegetation) NOT (marine OR aquatic) |
| Weeds | ((agri* OR agro* OR farm*) AND weed* AND (vegetation OR woody OR riparian OR plant*) AND biodiversity AND ecol* AND (effect OR impact) NOT review NOT wetland*) | ((agri* OR agro* OR farm*) AND weed* AND (vegetation OR woody OR riparian OR plant*) AND biodiversity AND ecol* AND (effect OR impact) NOT review NOT wetland*) |
| Abiotic | (agr* OR farm*) AND (vegetation OR non-crop) AND process AND function* AND eco* AND (abiotic OR decomposition OR sequestration OR carbon OR nutrient* OR respir* OR quality OR flamm* OR phyto* OR soil*) AND NOT (review OR wetland) | (agr* OR farm*) AND (vegetation OR non-crop) AND process AND function* AND eco* AND (abiotic OR decomposition OR sequestration OR carbon OR nutrient* OR respir* OR quality OR flamm* OR phyto* OR soil*) AND NOT (review OR wetland) |

**Table S2:** Descriptions of the non-production vegetation elements encountered in this review.

| Non-production<br>vegetation element | Description |
| --- | --- |
| Mixed | A mix of different non-production vegetation element types |
| Reserve | A large area of native vegetation outside of agricultural production |
| Corridor | A linear non-production vegetation element, specifically described as facilitating the movement of animals, pests or diseases within a landscape |
| Patch | A non-linear (round or rectangular) non-production vegetation element, greater than field scale |
| Riparian margins | Usually linear non-production vegetation element alongside rivers or creeks |
| Shelterbelts | Linear non-production vegetation elements planted on farms for shelter and shade |
| Hedgerows/borders | Linear non-production vegetation elements occurring between fields |
| Scattered individuals | Multiple individual trees within a farm |
| Plots | Small-scale (within field) experimental plantings |
| Single trees | An individual tree |
| Understorey | Non-production vegetation planted underneath orchard or other crop vegetation |

**Table S3.** Description of process categories collated from literature review.

| Process category | Combined category | Description |
| --- | --- | --- |
| Biocontrol | Biocontrol | Control of pests in production vegetation via predation that is not parasitoids and does not involve spillover from plantings or adjacent non-production elements |
| Biocontrol spillover | Spillover | Pest control via predation that is not parasitoids that involves predators moving from non-production elements into production vegetation and controlling pest prey |
| Parasitoid biocontrol | Biocontrol | Control of pests in production vegetation via predation by parasitoids |
| Parasitoid biocontrol spillover | Spillover | Pest control via parasitoid predation that involves parasitoids moving from non-production elements into production vegetation and controlling pest prey |
| Predation | Predation | Predation that does not involve control of pests in production vegetation (includes seed predation) |
| Predator spillover | Spillover | Predation of non-pest prey by animals that are not parasitoids that involves predators moving from non-production elements into production vegetation (predation may have been inferred or measured; includes seed predation) |
| Pollinator spillover | Spillover | Movement of animals that pollinate production vegetation from non-production elements into production vegetation (predation may have been inferred or measured) |
| Pest spillover | Spillover | Disservice: Movement of pest animals from non-production elements into production vegetation |
| Landscape dispersal | Dispersal | Dispersal at landscape scales between non-production or production elements that does not involve the use of corridors, nor explicitly tests matrix permeability (at larger scales than within a habitat or across habitat edges) |
| Corridor dispersal | Dispersal | Movement of animals through linear non-production vegetation elements at a landscape scale (not planted borders, hedgerows or strips within or next to fields; at larger scales than within a habitat or across habitat edges) |
| Matrix dispersal | Dispersal | Dispersal at landscape scales across inhospitable or impermeable landscape elements (at larger scales than within a habitat or across habitat edges) |
| Pest dispersal | Dispersal | Movement of pest animals at a landscape scale driven by the presence of non-production vegetation elements |
| Seed dispersal | Dispersal | Animal consumption of seeds or fruits without seed mortality |
| Habitat selection | Habitat selection | Choice and movement of animals between different landscape elements including either production and/ or non-production habitat types |
| Habitat provision | Habitat provision | Changes in animal population density driven by the presence of non-production vegetation elements |
| Pest competition | Competition | Disservice: Direct or indirect negative interactions between native and pest animal species driven by the presence of non-production vegetation elements |
| Pest habitat selection | Habitat selection | Disservice: Choice of pest animals between different landscape elements including either production and/ or non-production habitat types |
| Pest herbivory | Herbivory | Disservice: Damage of pest animals to production vegetation |
| Pest predation | Predation | Disservice: Predation by pest animals of native animals |

